# Sequence analysis of SARS-CoV-2 in nasopharyngeal samples from patients with COVID-19 illustrates population variation and diverse phenotypes, placing the in vitro growth properties of B.1.1.7 and B.1.351 lineage viruses in context

**DOI:** 10.1101/2021.03.30.437704

**Authors:** Tessa Prince, Xiaofeng Dong, Rebekah Penrice-Randal, Nadine Randle, Catherine Hartley, Hannah Goldswain, Benjamin Jones, Malcolm G. Semple, J. Kenneth Baillie, Peter J. M. Openshaw, Lance Turtle, ISARIC4C Investigators, Grant L. Hughes, Enyia R. Anderson, Edward I. Patterson, Julian Druce, Gavin Screaton, Miles W. Carroll, James P. Stewart, Julian A. Hiscox

## Abstract

New variants of SARS-CoV-2 are continuing to emerge and dominate the regional and global sequence landscapes. Several variants have been labelled as Variants of Concern (VOCs) because of perceptions or evidence that these may have a transmission advantage, increased risk of morbidly and/or mortality or immune evasion in the context of prior infection or vaccination. Placing the VOCs in context and also the underlying variability of SARS-CoV-2 is essential in understanding virus evolution and selection pressures. Sequences of SARS-CoV-2 in nasopharyngeal swabs from hospitalised patients in the UK were determined and virus isolated. The data indicated the virus existed as a population with a consensus level and non-synonymous changes at a minor variant. For example, viruses containing the nsp12 P323L variation from the Wuhan reference sequence, contained minor variants at the position including P and F and other amino acids. These populations were generally preserved when isolates were amplified in cell culture. In order to place VOCs B.1.1.7 (the UK ‘Kent’ variant) and B.1.351 (the ‘South African’ variant) in context their growth was compared to a spread of other clinical isolates. The data indicated that the growth in cell culture of the B.1.1.7 VOC was no different from other variants, suggesting that its apparent transmission advantage was not down to replicating more quickly. Growth of B.1.351 was towards the higher end of the variants. Overall, the study suggested that studying the biology of SARS-CoV-2 is complicated by population dynamics and that these need to be considered with new variants.

**Importance:** SARS-CoV-2 is the causative agent of COVID-19. The virus has spread across the planet causing a global pandemic. In common with other coronaviruses, SARS-CoV-2 genetic material (genomes) can become quite diverse as a consequence of replicating inside cells. This has given rise to multiple variants from the original virus that infected humans. These variants may have different properties and in the context of a widespread vaccination program may render vaccines less ineffective. Our research confirms the degree of genetic diversity of SARS-CoV-2 in patients. By isolating viruses from these patients, we show that there is a 100-fold range in growth of even normal variants. Interestingly, by comparing this to the pattern seen with two Variants of Concern (UK and South African variants), we show that at least in cells the ability of the B.1.1.7 variant to grow is not substantially different to many of the previous variants.

## Introduction

SARS-CoV-2 emerged late 2019 in Wuhan, China and causes COVID-19 (1). This can be a fatal infection with severe immunopathology in the respiratory system (2). The virus has since spread worldwide and resulted in more than 2.5 million deaths (3) placing large burdens on healthcare infrastructures and global economies. Several vaccines have been granted emergency licensure and these appear to be driving down cases in countries with large scale vaccine roll outs. However, multiple variants have been identified worldwide and these have the potential for vaccine evasion and immune escape, leading to the label of Variants of Concern (VOCs).

SARS-CoV-2 has a single stranded positive sense RNA genome about 30kb in length. The first two thirds of the genome is translated to give the viral non-structural proteins (NSP1-16), which includes the viral RNA dependent RNA polymerase (NSP12). Several viral RNA synthesis processes occur during infection including replication of the genome and transcription of a nested set of subgenomic mRNAs (sgmRNAs). This latter process requires discontinuous transcription during negative strand synthesis (4). As a natural consequence, coronaviruses have high levels of recombination. This can result in both deletions and insertions and template switching as well as the formation of defective RNAs. An example of this is the probable insertion of the furin cleavage site in the spike glycoprotein (5). Although SARS-CoV-2 and other coronaviruses have some type of proof-reading capability (6), this is generally thought to help maintain their large genomes, without entering error catastrophe. Otherwise the accumulation of deleterious mutations would result in a rapid loss of fitness and extinction of a viral population (7). Additionally, potential genome modifications can result from nucleotide changes through the action of cellular proteins involved in RNA processing (8). SARS-CoV-2 accumulates mutations at roughly the same frequency as Ebola virus (9). These drivers of genetic diversity and the numbers of people infected has led to multiple lineages and variants of SARS-CoV-2 being identified worldwide.

The sgmRNAs encode for the main structural proteins, including the envelope protein (E) protein, the membrane (M) protein, the nucleocapsid (N) protein and the spike (S) glycoprotein. The S protein is a component of the enveloped virion and interacts with the angiotensin converting enzyme-2 receptor (ACE-2) found on human cells. The S protein is also the major source of neutralising epitopes and therefore under selection pressure in coronaviruses (and SARS-CoV-2). Other viral proteins are involved in modulating the innate immune response.

Many variations in the coronavirus genome occur in the S gene (10–12) and this also has been identified for SARS-CoV-2. For example, the D614G substitution in the SARS-CoV-2 S protein, which emerged by March 2020, demonstrated improved transmissibility compared to Wuhan variants, and proceeded to dominate worldwide subsequently. This mutation is most often accompanied with another amino acid substitution in NSP12, P323L (13). In September 2020, a variant of concern, VOC 202012/01 (B.1.1.7 lineage) was detected in Kent in the UK which possessed 23 mutations distinct from the Wuhan reference sequence, including the N501Y substitution in the receptor binding domain of the S protein. This may increase the affinity of spike protein to ACE-2 receptor (14). Initial data suggested this variant could be related to an increased risk of hospitilisation and death (15). The variant has now spread to several countries and modelling studies have suggested increased transmissibility (16). Preliminary experiments in hamsters have identified increased viral shedding compared to the D614G variant (17). However, *in vitro* studies suggest that the B.1.1.7 VOC does not have any replicative advantage in primary airway epithelial cells (18).

As variants are likely to continue to emerge on a background of incomplete vaccination globally, understanding the significance of such variants both *in vitro* and *in vivo* is important to provide biological mechanistic data rather than rely on *in silico* modelling to determine their potential threat to vaccines or transmission advantage. To investigate the genetic and phenotypic diversity of SARS-CoV-2 in patients and in the context of the emergence of the B.1.1.7 and B.1.351 lineage viruses and concerns around potential higher viral loads, the growth of these viruses was bench marked against the Victoria isolate and clinical isolates from other samples taken during the outbreak.

## Results

Although consensus genomes for SARS-CoV-2 are reported on global databases from the sequencing of clinical specimens, in reality the virus will exist as a population within an individual and may also include defective RNAs. Likewise, in some pipelines, viral genomes or variants containing out of place stop codons within ORFs will not be returned as consensus even though they may be dominant. In this case, at a minor variant level, which could represent 49% of other genomes within the same individual, the wildtype protein may be expressed, and counterbalance any aberrantly functioning proteins. To investigate the sequence diversity of SARS-CoV-2 within a patient and to compare the growth of these viral populations to recent VOCs, nasopharyngeal swabs were taken from patients with COVID-19, sequenced and the genotypes and variants of isolated viruses and their growth properties compared in cell culture (Figure 1).

**Figure 1.**
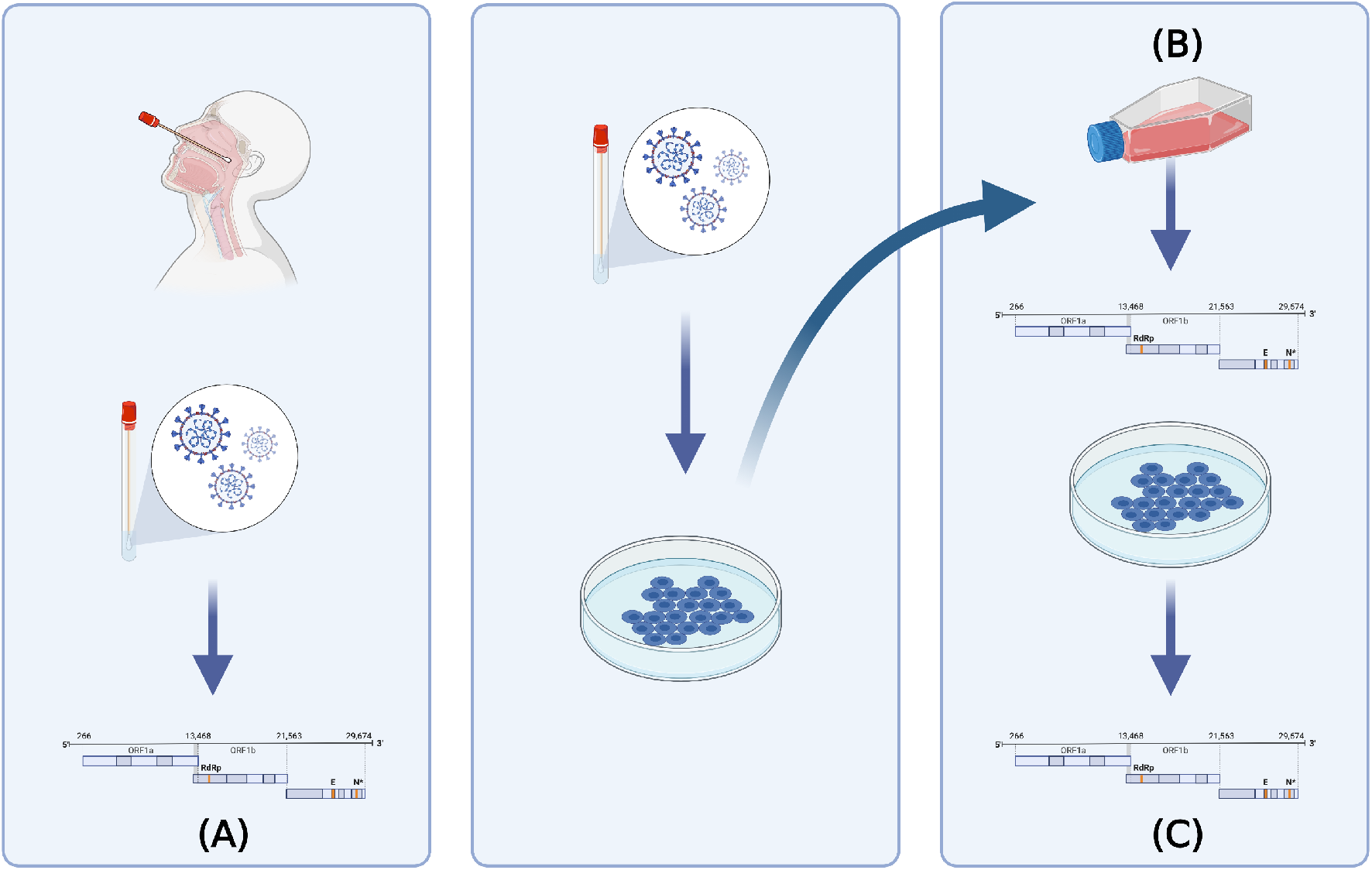
Testing strategy. (A) Nasopharyngeal swabs from patients with COVID-19 recruited to the ISARIC-4C study were sequenced using an amplicon based approach on the Oxford Nanopore MinION (P0). Virus was isolated from the same nasopharyngeal swabs(P1). (B) Viral isolates from the ISARIC-4C study, B.1.1.7, B.1.351 and Victoria isolates were grown up into stocks which were then sequenced. (C) Viral stocks were titrated and used to infected hACE2-A549 cells, and 72-hour post infection supernatants were sequenced.

### Sequence variation of SARS-CoV-2 in clinical swabs compared to Wuhan reference strain

Nine swabs representing different time points in the outbreak in the UK contained recoverable virus that could be isolated and grown. The virus population in these swabs was sequenced and both consensus genomes and minor variants determined. Consensus sequence variation was compared to the reference genome (NC_045512; Wuhan-Hu-1) to see how far the isolates had diverged and with minor variants listed for the secondary and tertiary positions (Supplementary Table 1). Most viruses demonstrated a few amino acid variations compared to the reference sequence. For example, SCV2-006, a lineage B virus sequenced from the swab of a patient from the Diamond Princess cruise ship (February 2020) had only one substitution present, R203K in the N protein (Figure 2). In comparison, sequence analysis of SCV2-009, a virus isolated from a swab sampled from a patient in the UK in March 2020 (Figure 2) now possessed the D614G and P323L substitutions in the spike glycoprotein and NSP12, respectively. These are in contrast to the B.1.1.7 variant, which emerged later in 2020 and is characterised by the presence of 23 amino acid differences from the reference genome. Analysis of the virus population present in the nasopharyngeal isolate of SCV2-009 illustrated the diversity associated with the virus. For example, taking the P323L substitution in NSP12, out of an amino acid coverage of 202, 170 amino acids mapped to L, 12 to P and 9 to F. For the D614G substitution in the spike glycoprotein, out of an amino acid coverage of 3452, 3360 amino acids mapped to G, 24 to S and 21 to V. This general pattern is reflected in other clinical isolates. For example, in isolate SCV2-010, in NSP12, out of an amino acid coverage of 285, 273 mapped to L, 50 to I and 3 to P. In isolate SCV2-008, in NSP12, out of an amino acid coverage of 153, 130 mapped to L, 9 to P and 7 to F. This suggests, for NSP12, that at the minor variant level the reference sequence amino acid is still present, but other amino acids such as F may be common (Supplementary Table 1), and subject to selection pressure. In some clinical swabs, for example in N at position 204, the second most common feature is a stop codon. SCV2-011 and SCV2-018 were variants isolated from clinical swabs taken from the same patient but three days apart, these did not vary at the consensus between each other, but did at the minor variant level. SCV2-007 and SCV2-017 were also variants isolated from clinical swabs taken from the same patient but three days apart and did not vary at the consensus between each other in swabs, but did at the minor variant level.

**Figure 2.**
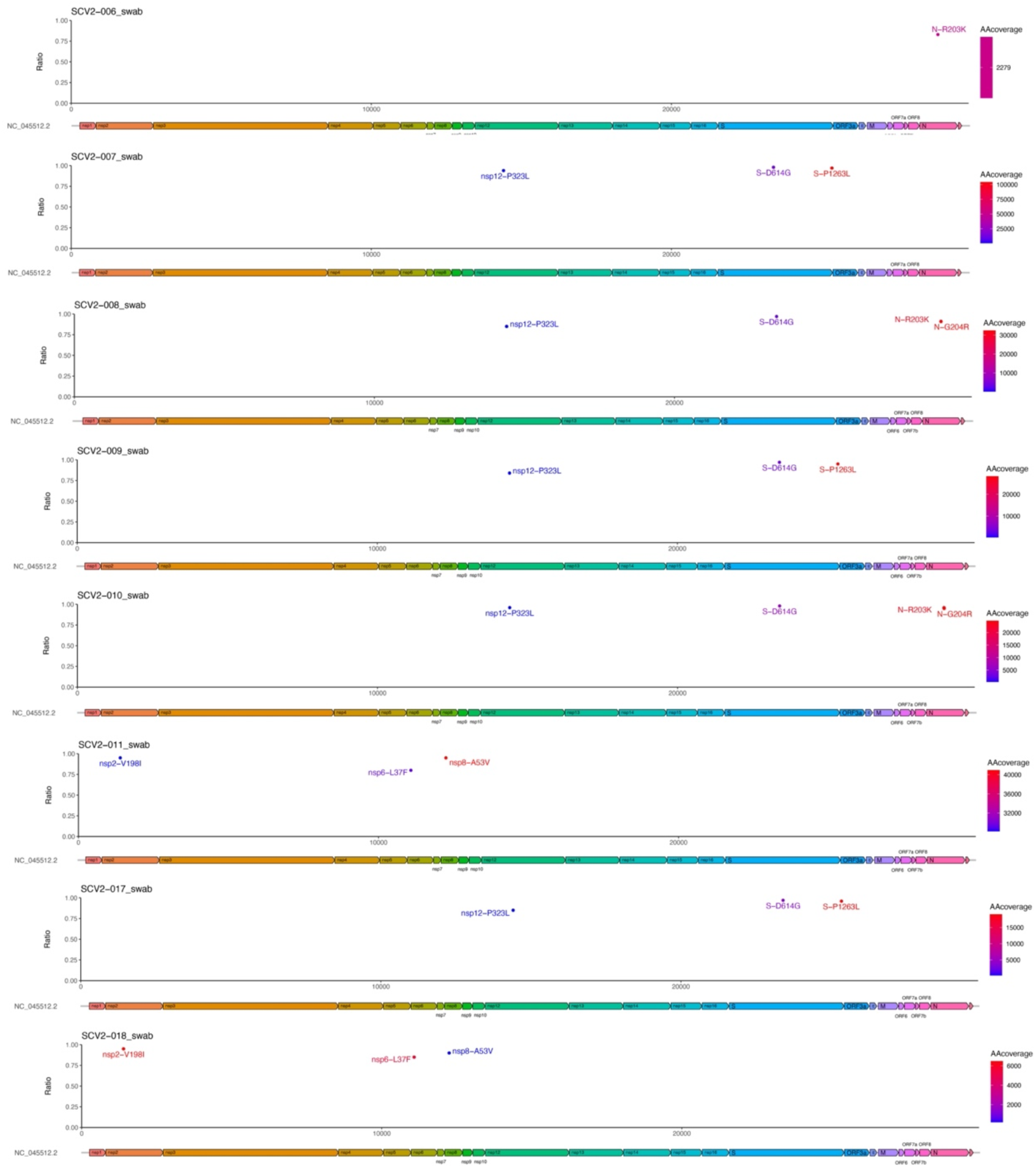
(A) Comparison of the consensus sequence SARS-CoV-2 in ISARIC4C swabs collected from patients in hospital with COVID-19. Variations from the Wuhan reference sequence are plotted with the location on the viral genome against the percentage (ratio) of reads where the variation is observed. Variants are coloured to demonstrate the number of reads achieved (>10).

### Comparison of sequence variation in stocks and after 72 hours in hACE2-A549 cells

In order to assess the biology of the viruses isolated from the clinical swabs and compare their growth to B.1.1.7 and B.1.351, sufficient stocks had to be grown. To isolate SARS-CoV-2 from the clinical swabs, the nasopharyngeal sample was filtered and placed on VeroE6 cells with antibiotics and antifungals until CPE was observed. The supernatant was collected from these cells to generate sufficient stocks for infectivity assays and comparisons.

Growing virus for stocks may have introduced or selected for specific variants. One of these, that has been characterised for SARS-CoV-2, is a deletion of the furin cleavage site in the spike glycoprotein when grown in Vero E6 cell (19). Therefore, viral stocks were sequenced to ensure they did not possess the deletion and to determine if variation occurred compared to when the virus was sequenced directly from clinical swabs. Comparator viruses of known provenance were obtained from collaborators. The comparator viruses were the B.1.1.7 (‘Kent’ UK VOC) virus (termed SCV2-019 in this study) obtained at P4, the SARS-CoV-2/Victoria/01/2020 (an isolate from Australia) obtained at P3 (termed SCV2-021), and the B.1.351 virus (‘South African’ VOC) (termed SCV2-022 in this study). These were grown in Vero/hSLAMS as a precaution to prevent selection for the furin deletion. These were also sequenced to ensure they had the variant defining mutations present. For these three comparator viruses, the sequencing showed at the consensus level the furin cleavage site was intact and the other defining variations separating these variants from the Wuhan reference sequence were present (Figure 3).

**Figure 3.**
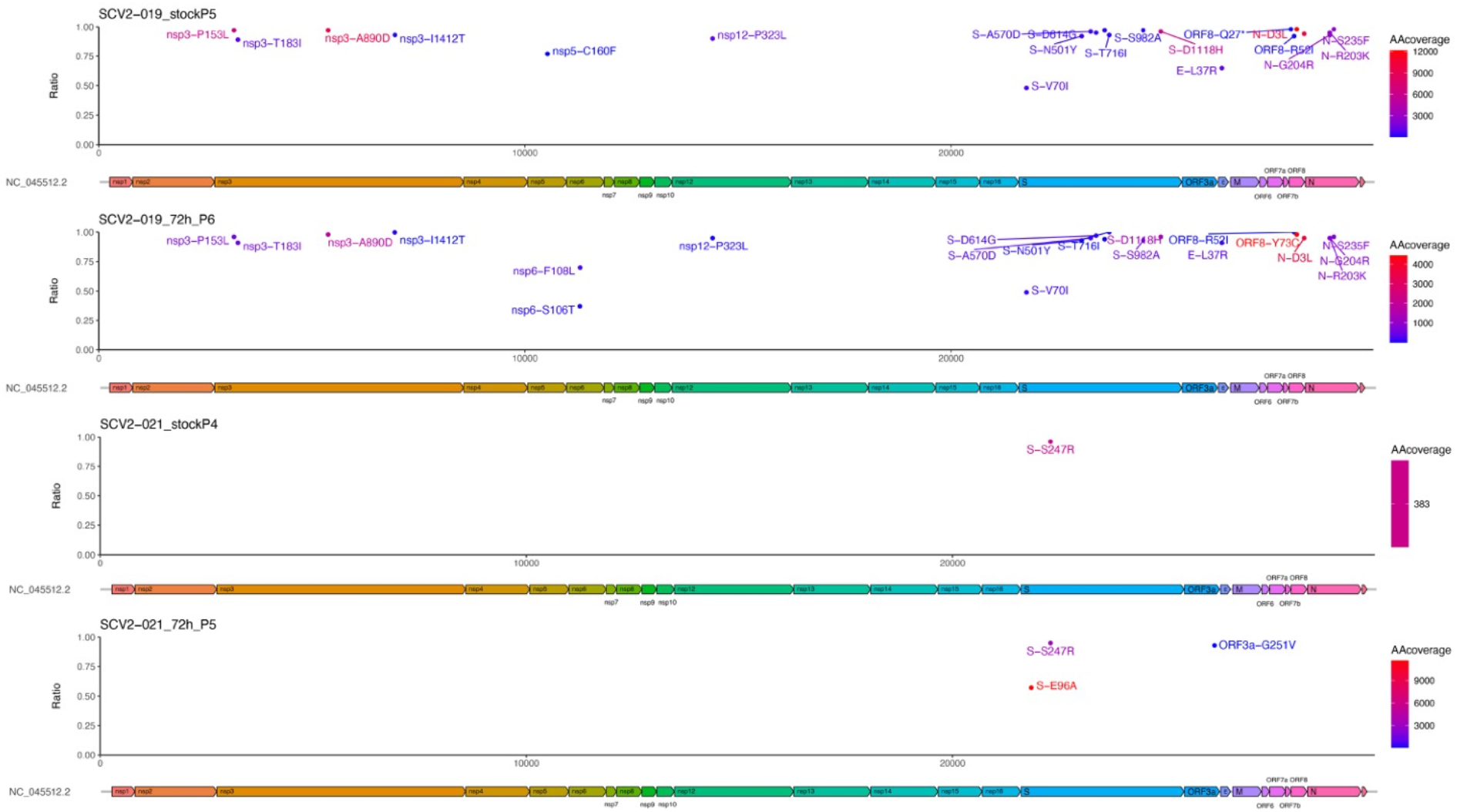
Comparison of the UK ‘Kent’ VOC (SCV2-019) and the Australian Victoria isolate (SCV2-021) with the Wuhan reference sequence. Variations from the Wuhan reference sequence are plotted with the location on the viral genome against the percentage (ratio) of reads where the variation is observed. Variants are coloured to demonstrate the number of reads achieved (> 10).

Analysis of the genome diversity between viruses sequenced in swabs from patients and the virus stock used to infect cells indicated that most consensus variations from the Wuhan reference sequence were still present (Supplementary Table 1). The minor variants at selected positions were also still present. For example, in the stock preparation for SCV2-009, at position 323 in NSP12, this was read with an amino acid depth of 517. The L was present at a depth of 499, P with a depth of 6 and F with a depth of 5, indicating that the consensus level amino acid was still present with P and F at a minor level. For some stock viruses, variation from the reference sequence was lost during preparation of the stock virus. The growth of these viruses from the stocks was compared to a B.1.1.7. and a B.1.351 lineage virus and SARS-CoV-2/Victoria/01/2020, obtained from near the start of the COVID-19 pandemic.

### Growth comparison of different SARS-CoV-2 variants to Variants of Concern (VOCs)

In order to identify whether the B.1.1.7 (SCV2-019) and B.1.351 (SCV2-022) displayed a growth advantage over less recent strains of the virus, three different cell lines were infected with the viruses at an MOI of 0.01 over the course of 72 hours and the resultant supernatants, at 24, 48 and 72 hours titrated by plaque assay on Vero E6 cells. The three different cell lines were Vero E6 (commonly used to grow viral stocks and initial isolates from clinical samples), Vero/hSLAM (reported to prevent deletion of the furin cleavage site in the spike glycoprotein) and hACE2-A549 cells. This latter cell line is based on A549 cells, which are respiratory epithelium in origin, commonly used to study respiratory viruses in cell culture but overexpress the ACE2 protein. A549 cells mount an interferon response to virus infection.

In Vero E6 cells, eleven SARS-CoV-2 variants followed a similar pattern of growth, with the exception of SCV2-021 (SARS-CoV-2/Victoria/01/2020) which grew at significantly reduced levels compared to other variants by 72 hours post infection (mean 5.7 × 10^4^ PFU/ml, p=0.006) (Figure 4A). A similar pattern of growth was observed in all twelve variants in Vero/hSLAM cells, but there was no significant difference (p>0.05) in the titres of any of the viruses produced by 72 hours post infection (Figure 4B). In contrast, in hACE2-A549 cells, there was more heterogeneity observed between variants, with the range of viral titres being much lower (7.9 × 10^1^ −3.01 × 10^4^ PFU/ml) than that observed in Vero cells at 72 hours post infection. The B.1.1.7 variant (SCV2-019) had the lowest titre at 24 hours post-infection in growth assays in hACE2-A549 cells, before growing to reach a final titre 2.71 × 10^3^ PFU/ml at 72 hours post-infection (Figure 4C). We observed that between variants at the growth extremes at 72 hrs post-infection in hACE2-A549 cells there was an approximately <2 log difference between titres of SCV2-007 and SCV2-018, despite the same amount of virus being used as the inoculum (MOI=0.01). For the VOC B.1.1.7 the growth at 72 hrs post-infection in hACE2-A549 cells was in the middle of the other variants tested (2.71 × 10^3^ PFU/ml) while the VOC B.1.351 had the second highest final titre of 2.62 × 10^4^ PFU/ml. (Figure 4C). In addition, we note that the B.1.351 lineage variant had the highest viral titres in both VeroE6 and Vero/hSLAMs at 24 and 48 hrs post-infection.

**Figure 4.**
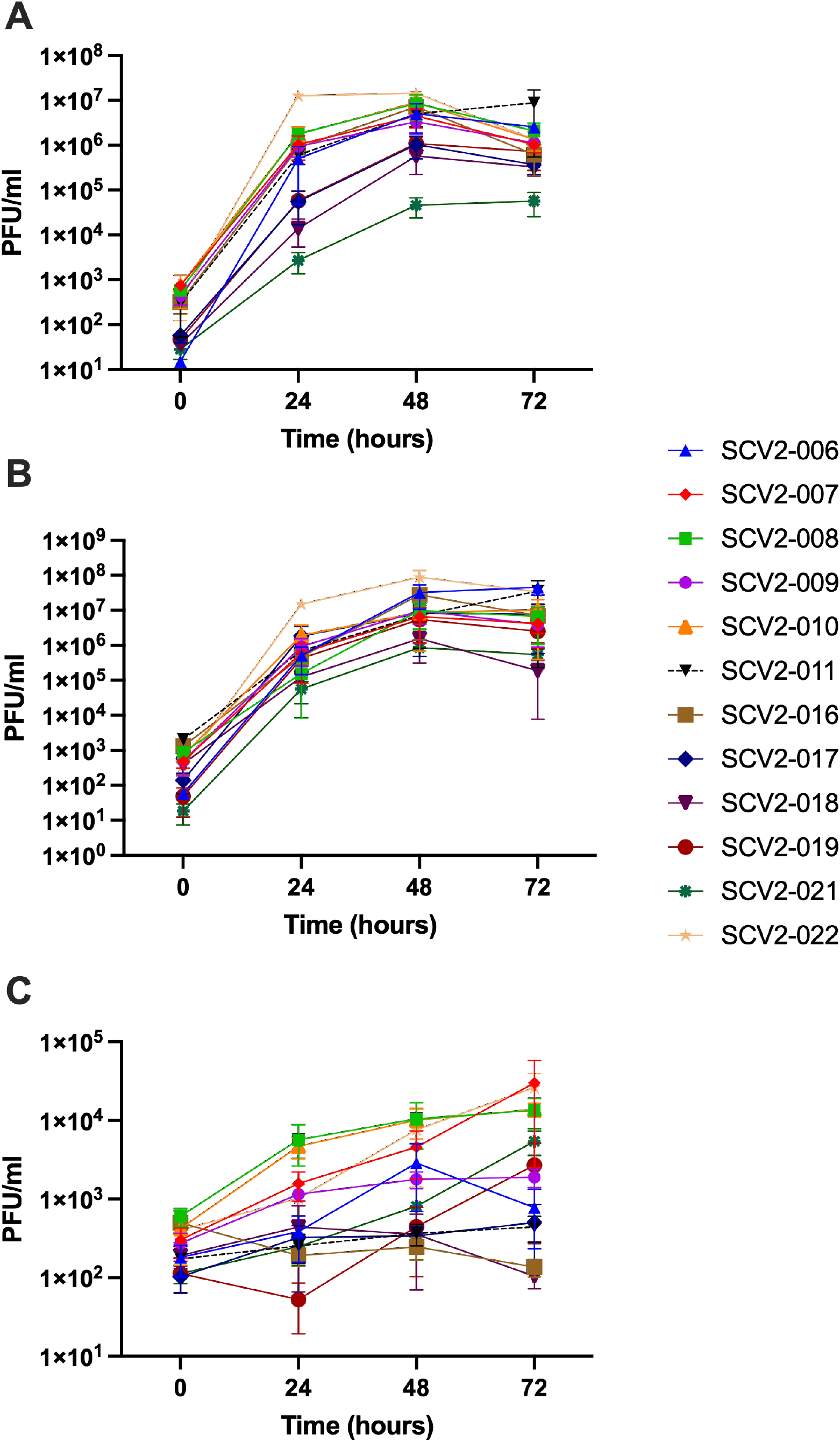
Growth over time of 11 different viral isolates in three different cell lines compared to the Variants of Concern SCV2-019 (UK ‘Kent’ VOC) and SCV2-022 (‘South African’ VOC). Comparison viruses included the Australian Victoria variant (SCV2-021). (A) Growth of viruses in plaque-forming units (PFU) per ml over times in Vero E6 African green monkey kidney cells. (B) Growth of viruses in Vero cells expressing the human signalling lymphocytic activation module (SLAM) gene (Vero/hSLAM). (C) Growth of viruses in human ACE-2 expressing A549 cells (hACE2-A549). All experiments were repeated in triplicate using supernatant from 6 wells (n=3).

Comparing viruses grown from the same patient but sampled three days apart (SCV2-007 and SCV2-017, and SCV2-011 and SCV2-018 at day one and day three respectively, showed differences in their growth in hACE2-A549 cells. (Supplementary Table 2). There was a 2-log difference in the growth of SCV2-007 and SCV2-017, while there was little difference in the growth of SCV2-011 and SCV2-018.

### The phenotype of the variants differed widely between cell lines, displaying mixed plaque morphology and growth characteristics

The phenotype of the plaques formed by each virus stock was observed in the three different cell lines used at 72 hours post-infection. The appearance of the plaques from the variants differed (Figure 5). SCV2-006, SCV2-011, SCV2-016, SCV2-018 and SCV2-022 had a larger plaque phenotype after growth in Vero E6 cells, compared with SCV2-019 (B.1.1.7) and SCV2-021 (SARS-CoV-2/Victoria/01/2020). Equally, some variants displayed a mixed phenotype of both large and small plaques in Vero E6 cells, as seen for SCV2-011, suggesting mixed viral species were present (Figure 5). After growth in Vero/hSLAMS, SCV2-011 and SCV2-018 showed a mixed phenotype after plaque assay. SCV2-019 (B.1.1.7) and SCV2-021 had the smallest plaque phenotypes. After growth in hACE2-A549 cells, SCV2-006 and SCV2-022 (B.1.351) had the largest plaque phenotypes, while SCV2-021 had the smallest plaque phenotype. SCV2-006 and SCV2-016 had mixed morphology of both large and small plaques. This illustrates the potential diversity with a viral population.

**Figure 5.**
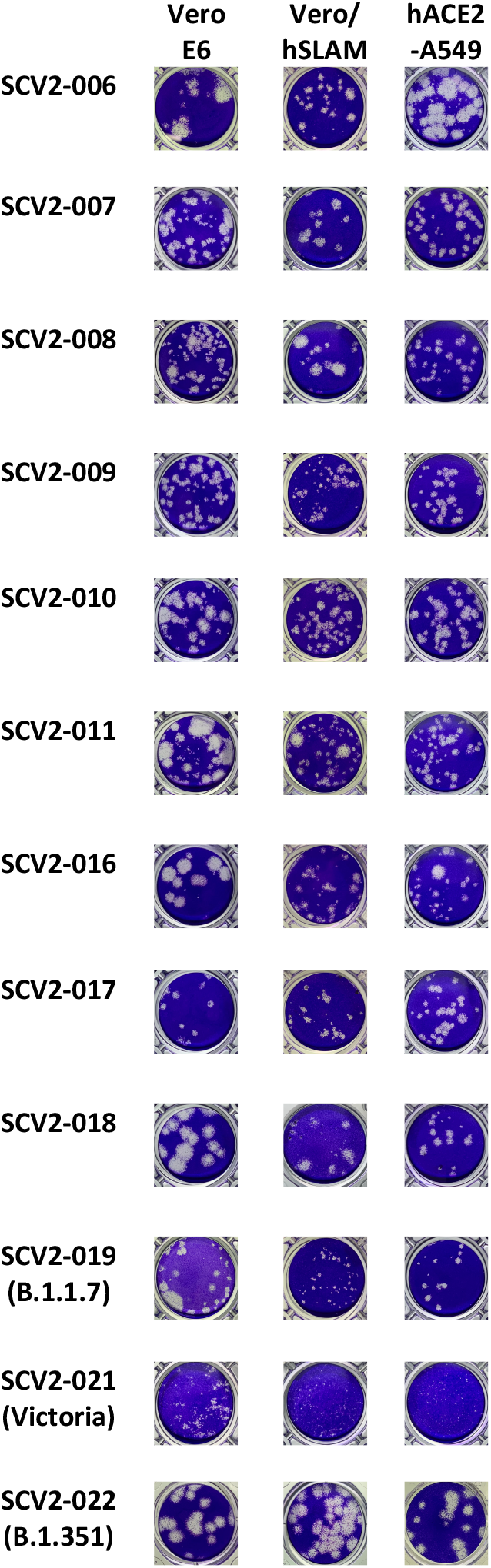
Phenotypic appearance of plaque assays from variants grown in three different cell lines; (i) Vero E6, (ii) Vero/hSLAM and (iii) hACE2-A549 cells. Plaque assays were performed on VeroE6 cells. Variants of Concern are SCV2-019 (UK ‘Kent’ VOC) and SCV2-022 (‘South African’ VOC). Comparison viruses include the Australian Victoria variant (SCV2-021).

### Genetic diversity of variants after passage in the three different cell types

We hypothesised that differences in the phenotypic appearance of viruses and their reproduction might reflect the presence of minor variants and stop codons in their underlying sequences and this was investigated at 72 hrs post-infection in hACE2-A549 cells. All variants have a consensus level genome but also minor variants. In SCV2-016, at 72 hrs post-infection there was a stop codon at consensus level in ORF3A that was also present in the viral stock (Supplementary Table 1). In SCV2-019 (the B.1.1.7. lineage virus) there was a stop codon in ORF8 at the consensus level at 72 hr post-infection, which was also present in the stock (Supplementary Table 1). We note that both of these were low read depth, and other amino acids were present at the minor variant level. Stop codons were also present in the variants at a minor variant level (Supplementary Table 1).

## Discussion

Sequence analysis of SARS-CoV-2 in clinical swabs from patients revealed a heterogenous and diverse population from the Wuhan reference sequence. At the minor variant level, a number of variants had genomes which contained premature stop codons. Examples of SARS-CoV-2 genomes encoding non-functioning proteins have been previously identified in the human population. For example, a cohort of patients in Singapore were identified with a deletion in ORF8 that was associated with a milder infection (20), although the variant disappeared either through control measures or lack of fitness. This potential disconnect is not restricted to SARS-CoV-2. The balance between consensus and minor variants and the presence of stop codons in virus populations within individual patients has been shown to influence the activity of the Ebola virus RNA dependent RNA polymerase and correlate with outcome in patients with Ebola virus disease (21). Within an individual person with SARS-CoV-2, these mixtures of functioning and presumably non-functioning viral proteins will potentially influence viral load.

Recent VOCs include SARS-CoV-2 variants from Nigeria (B.1.525). One of the differences in this variant from the Wuhan consensus sequence is P323F in NSP12. This variation was also identified in a cluster of patients in Northern Nevada in the USA (22). The analysis of variants in this study, isolated earlier in 2020, indicated that an F at position 323 existed at a minor variant level. Therefore, we hypothesize that if an F at this position was advantageous (e.g. altered RdRp activity) then the variant would have been selected during passage. However, this was not the case, and therefore we speculate that the emergence of an F at position 323 in NSP12 may be through founder effect.

The growth of different variants, and a variant from near the start of the COVID-19 pandemic, were compared in three different cell lines to the growth of two VOCs and the Australia Victoria variant. These VOCs were from the B.1.1.7 and B.1.351 lineages and represent viruses that have an apparent transmission advantage in the general population and/or may be less refractive to currently approved vaccines. There was an approximately 2 log difference in growth at 72 hrs in the hACE2-A549 cells by the different variants. The growth of VOCs in this cell line were within these limits. Extrapolating this observation to the perceived transmission advantage of B.1.1.7 in the human population, would suggest this is not down to the VOC growing to higher titres in cells in vivo compared with other variants, though we acknowledge that out *in vitro* experiments may not correlate exactly with growth rates *in vivo*. With B.1.1.7, the same heterogenous patterns of disease in humans as other variants, although we note that B.1.1.7 has been associated with a decrease in Ct values from nasopharyngeal swabs and also an increase in mortality in some populations. In contrast, different variants of the coronavirus infectious bronchitis (IBV) can individually cause different spectrums of organ specific disease, and therefore current variations in the genome of SARS-CoV-2 may not automatically equate to radically different disease as observed with IBV. Due to the promiscuous nature of coronavirus RNA synthesis, variants have and will occur all of the time. This emphasises the need for genotype to phenotype studies to place newly emerged variants that have perceived differences in context.

Comparison of viruses isolated from the same patients at different time points revealed intriguing differences. While the viruses SCV2-007 and SCV2-017 differed by 2-logs in hACE2-A549 cells at 72 hours, there was little difference observed between the twin viruses SCV2-011 and SCV2-018. Notably, the SCV2-017 virus had picked up an additional mutation at 72 hours post-infection – a change from A in the reference genome at position 1120 in NSP3 to a V. It is possible this may be responsible for the difference in growth of this virus to its founder, SCV2-007, and could reflect viral adaptation to the immune response in this individual over the course of infection (Supplementary table 2).

The analysis of virus in the nasopharyngeal swabs clearly paints a picture of a diverse population of SARS-CoV-2. When studying isolates, even when grown in cell culture, that population still continues. Thus, whilst lineage defining variations are present at a consensus level, minor variants are present underneath that may have an impact on biology. This would suggest that the study of specific genotypes requires either plaque purification or reverse genetics. However, the study suggests that the viral population (consensus and minor variants) should be taken into account when studying the transmission of SARS-CoV-2.

## Methods

### Cells

African green monkey kidney C1008 (Vero E6) cells (Public Health England, PHE) were cultured in Dulbecco’s minimal essential medium (DMEM) (Sigma) with 10% foetal bovine serum (FBS) (Sigma) and 0.05mg/ml gentamicin at 37°C/5% CO_2_. Vero/hSLAM cells (PHE) were grown in DMEM with 10% FBS and 0.05mg/ml gentamicin (Merck) with the addition of 0.4mg/ml Geneticin (G418; Thermofisher) at 37°C/5% CO_2_. Human ACE2-A549 (hACE2-A549), a lung epithelial cell line which overexpresses the ACE-2 receptor, were the kind gift of Oliver Schwartz (23) and were cultured in DMEM with 10% FBS and 0.05mg/ml gentamicin with the addition of 10μg/ml Blasticidin (Invitrogen). Only passage 3-10 cultures were used for experiments.

### Virus isolation

The SARS-CoV-2/human/Liverpool/REMRQ0001/2020 isolate (Genbank ID MW041156.1), was used at passage 3. The fourth passage of virus (here named SCV2-006_stockP4) was cultured in Vero E6 cells with DMEM containing 4% FBS and 0.05mg/ml gentamicin at 37°C/5% CO_2_ and harvested 48 hours post inoculation. Virus stocks were aliquoted and stored at −80°C.

Viruses named SCV2-007 to SCV2-018 were grown from nasopharyngeal swabs of patients using the following method. One hundred microlitres of viral transport media from the swab was mixed with 100μl DMEM with 4% FBS, 0.05mg/ml gentamicin, 25μg/ml plasmocin (Invivogen) and 2.5μg/ml amphotericin B (Merck). These were then filtered using ultrapure MC 0.22μm filters (Merck) and the filtrate placed onto cells in a 24 well plate of Vero E6 cells for 1 hour. After one hour, the media was topped up with DMEM (2% FBS, 0.05 mg/ml gentamicin, 25μg/ml plasmocin, 2.5 μg/ml amphotericin B). Cells were observed daily for cytopathic effect (CPE) and the cell supernatant harvested once CPE was evident. This provided the first passage virus. Stocks of these were then grown in Vero E6 as described above and frozen down in aliquots at −80°C and named SCV2-007 to SCV2-018_stockP2.

The B.1.1.7 and B.1.351 isolates were used at passage 4. The fifth passage (here named SCV2-019_stockP5 and SCV2-022_stockP5) were cultured in Vero/hSLAM cells with DMEM containing 4% FBS, 0.05mg/ml gentamicin and 0.4mg/ml geneticin and harvested 72 hours post inoculation. Virus stocks were aliquoted and stored at −80°C. SARS-CoV-2 Victoria/01/2020 was passaged three times in Vero/hSLAM cells. The fourth passage stock (here named SCV2-021_stockP4) was cultured in Vero/hSLAM cells DMEM containing 4% FBS, 0.05mg/ml gentamicin and 0.4mg/ml geneticin and harvested 72 hours post inoculation. Virus stocks were aliquoted and stored at −80°C (Supplementary table 2).

### Virus titration

Viral titres of stocks were calculated using plaque assays. Briefly, confluent 24-well plates of Vero E6 cells were inoculated with serial ten-fold dilutions of the stocks in duplicate for one hour at 37°C/5% CO_2_. Plates were overlaid with DMEM containing 2% FBS, 0.05mg/ml gentamicin and 2% low melting point agarose (Lonza) and incubated at 37°C/5% CO_2_ for 72 hours. Plates were fixed using 10% formalin, the overlay removed, and plates stained using crystal violet solution (Sigma). Virus titre was measured in plaque forming units per ml (PFU/ml).

### Virus growth kinetics

Vero E6, Vero/hSLAM and hACE2-A549 cells were grown in 96 well plates for viral growth kinetic experiments. For infection, media was removed from plates and virus inoculum added at an MOI of 0.01 in DMEM containing 2% FBS, 0.05mg/ml gentamicin and the respective selective antibiotics for each cell line (6 wells per timepoint). Plates were incubated at 37°C/5% CO_2_ for one hour. The inoculum was removed, and cells were washed once with PBS (Sigma). The respective media with 2% FBS (100μl) was added to each well. The cell supernatant was removed from wells and combined (0hrs post infection) and plates incubated further. Supernatants were likewise removed at 24, 48 and 72 hours post infection. Approximately 250μl of the supernatants were aliquoted directly into tubes containing 750μl Trizol LS (Fisher) to inactivate the virus. All supernatants and inactivated supernatants were stored at −80°C until viral titration and RNA extraction could be performed. All infections were performed at least three times in independent experiments.

### RNA extraction and amplification of viral nucleic acids

RNA from clinical samples was extracted and DNase treated as described previously. Samples from patients were sequenced using the RLSA approach (24). RNA from viral stocks and from 72-hour post infection cultures were sequenced by Oxford Nanopore long read length sequencing on flow cells run on MinION or GridION.

### Nanopore sequencing

Sequencing libraries for amplicons generated by RSLA (24) or ARTIC were prepared following the ‘PCR tiling of SARS-CoV-2 virus with Native Barcoding’ protocol provided by Oxford Nanopore Technologies using LSK109 and EXP-NBD104/114.

### Variant calling

The artic-ncov2019 pipeline v1.2.1 (https://artic.network/ncov-2019/ncov2019-bioinformatics-sop.html) was used to filter the passed Fastq files produced by Nanopore sequencing with lengths between 800 and 1600 for RSLA, and 400 and 700 for ARTIC. This pipeline was then used to map the filtered reads on the reference SARS-CoV-2 genome (NC_045512.2) by minimap2 and assigned each read alignment to a derived amplicon and excluded primer sequences based on the RSLA and ARTIC V3 primer schemes in the bam files. These bam files were further analysed using DiversiTools (http://josephhughes.github.io/btctools/) with the “-orfs” function to generate the ratio of amino acid change in the reads and coverage at each site of protein in comparison to the reference SARS-CoV-2 genome (NC_045512.2). The amino acids with highest ratio and coverage > 10 were used to assemble the consensus protein sequences.

### Statistics

Viral titre data was log transformed and one-way ANOVAs performed with post-hoc Bonferroni tests performed to determine if any significant difference at T=72 hours post infection occurred between the SCV2-019 (B.1.1.7) and other viruses in different cell lines.

### Ethics and clinical information

The patients from which the virus samples were obtained gave informed consent and were recruited under the International Severe Acute Respiratory and emerging Infection Consortium (ISARIC) WHO Clinical Characterisation Protocol CCP. Ethical approval for data collection and analysis by ISARIC4C was given by the South Central-Oxford C Research Ethics Committee in England (reference 13/SC/0149), and by the Scotland A Research Ethics Committee (reference 20/SS/0028). Samples were use with consent from patients or consultees. The ISARIC WHO CCP-UK study was registered at https://www.isrctn.com/ISRCTN66726260 and designated an Urgent Public Health Research Study by NIHR. Protocol, patient information sheets, consents, case report forms and process of data and sample access request are available at https://ISARIC4C.net.

### Biosafety

All work was performed in accordance with risk assessments and standard operating procedures approved by the University of Liverpool Biohazards Sub-Committee and by the UK Health and Safety Executive. Work with SARS-CoV-2 was performed at containment level 3 by personnel equipped with respirator airstream units with filtered air supply.

## Author Contributions

TP, XD, RP-R, NR, CH, HG, BJ, JD, GLH, GS, ERA and EIP performed the experiments, sequencing, bioinformatics and isolated virus. Data was analysed by TP, XD, RP-R and JAH. MWC, LT, JPS and JAH supervised the project. MGS, JKB and PJMO established the ISARIC4C consortium that was used to obtained some of the UK clinical isolates used in the study. TP, XD, MWC and JAH wrote the manuscript, all authors provided editing and final approval.

## Acknowledgements

Clinical isolates used in this study were gathered under the auspices of the ISARIC Coronavirus Clinical Characterisation Consortium (ISARIC4C) and processed at the University of Liverpool. We would like to acknowledge all members of the consortia. Consortium Lead Investigator: J. Kenneth Baillie; Chief Investigator: Malcolm G. Semple; Co-Lead Investigator: Peter J.M. Openshaw; ISARIC Clinical Coordinator: Gail Carson; Co-Investigators: Beatrice Alex, Benjamin Bach, Wendy S. Barclay, Debby Bogaert, Meera Chand, Graham S. Cooke, Annemarie B. Docherty, Jake Dunning, Ana da Silva Filipe, Tom Fletcher, Christopher A. Green, Ewen M. Harrison, Julian A. Hiscox, Antonia Ying Wai Ho, Peter W. Horby, Samreen Ijaz, Saye Khoo, Paul Klenerman, Andrew Law, Wei Shen Lim, Alexander J. Mentzer, Laura Merson, Alison M. Meynert, Mahdad Noursadeghi, Shona C. Moore, Massimo Palmarini, William A. Paxton, Georgios Pollakis, Nicholas Price, Andrew Rambaut, David L. Robertson, Clark D. Russell, Vanessa Sancho-Shimizu, Janet T. Scott, Thushan de Silva, Louise Sigfrid, Tom Solomon, Shiranee Sriskandan, David Stuart, Charlotte Summers, Richard S. Tedder, Emma C. Thomson, A.A. Roger Thompson, Ryan S. Thwaites, Lance C.W. Turtle, and Maria Zambon; Project Managers: Hayley Hardwick, Chloe Donohue, Ruth Lyons, Fiona Griffiths, and Wilna Oosthuyzen; Data Analysts: Lisa Norman, Riinu Pius, Tom M. Drake, Cameron J. Fairfield, Stephen Knight, Kenneth A. Mclean, Derek Murphy, and Catherine A. Shaw; Data and Information System Managers: Jo Dalton, James Lee, Daniel Plotkin, Michelle Girvan, Egle Saviciute, Stephanie Roberts, Janet Harrison, Laura Marsh, Marie Connor, Sophie Halpin, Clare Jackson, and Carrol Gamble; Data Integration and Presentation: Gary Leeming, Andrew Law, Murray Wham, Sara Clohisey, Ross Hendry, and James Scott-Brown; Material Management: William Greenhalf, Victoria Shaw, and Sarah McDonald; Patient Engagement: Seán Keating; Outbreak Laboratory Staff and Volunteers: Katie A. Ahmed, Jane A. Armstrong, Milton Ashworth, Innocent G. Asiimwe, Siddharth Bakshi, Samantha L. Barlow, Laura Booth, Benjamin Brennan, Katie Bullock, Benjamin W.A. Catterall, Jordan J. Clark, Emily A. Clarke, Sarah Cole, Louise Cooper, Helen Cox, Christopher Davis, Oslem Dincarslan, Chris Dunn, Philip Dyer, Angela Elliott, Anthony Evans, Lorna Finch, Lewis W.S. Fisher, Terry Foster, Isabel Garcia-Dorival, Willliam Greenhalf, Philip Gunning, Catherine Hartley, Antonia Ho, Rebecca L. Jensen, Christopher B. Jones, Trevor R. Jones, Shadia Khandaker, Katharine King, Robyn T. Kiy, Chrysa Koukorava, Annette Lake, Suzannah Lant, Diane Latawiec, L. Lavelle-Langham, Daniella Lefteri, Lauren Lett, Lucia A. Livoti, Maria Mancini, Sarah McDonald, Laurence McEvoy, John McLauchlan, Soeren Metelmann, Nahida S. Miah, Joanna Middleton, Joyce Mitchell, Shona C. Moore, Ellen G. Murphy, Rebekah Penrice-Randal, Jack Pilgrim, Tessa Prince, Will Reynolds, P. Matthew Ridley, Debby Sales, Victoria E. Shaw, Rebecca K. Shears, Benjamin Small, Krishanthi S. Subramaniam, Agnieska Szemiel, Aislynn Taggart, Jolanta Tanianis-Hughes, Jordan Thomas, Erwan Trochu, Libby van Tonder, Eve Wilcock, and J. Eunice Zhang; Local Principal Investigators: Kayode Adeniji, Daniel Agranoff, Ken Agwuh, Dhiraj Ail, Ana Alegria, Brian Angus, Abdul Ashish, Dougal Atkinson, Shahedal Bari, Gavin Barlow, Stella Barnass, Nicholas Barrett, Christopher Bassford, David Baxter, Michael Beadsworth, Jolanta Bernatoniene, John Berridge, Nicola Best, Pieter Bothma, David Brealey, Robin Brittain-Long, Naomi Bulteel, Tom Burden, Andrew Burtenshaw, Vikki Caruth, David Chadwick, Duncan Chambler, Nigel Chee, Jenny Child, Srikanth Chukkambotla, Tom Clark, Paul Collini, Catherine Cosgrove, Jason Cupitt, Maria-Teresa Cutino-Moguel, Paul Dark, Chris Dawson, Samir Dervisevic, Phil Donnison, Sam Douthwaite, Ingrid DuRand, Ahilanadan Dushianthan, Tristan Dyer, Cariad Evans, Chi Eziefula, Chrisopher Fegan, Adam Finn, Duncan Fullerton, Sanjeev Garg, Sanjeev Garg, Atul Garg, Effrossyni Gkrania-Klotsas, Jo Godden, Arthur Goldsmith, Clive Graham, Elaine Hardy, Stuart Hartshorn, Daniel Harvey, Peter Havalda, Daniel B. Hawcutt, Maria Hobrok, Luke Hodgson, Anil Hormis, Michael Jacobs, Susan Jain, Paul Jennings, Agilan Kaliappan, Vidya Kasipandian, Stephen Kegg, Michael Kelsey, Jason Kendall, Caroline Kerrison, Ian Kerslake, Oliver Koch, Gouri Koduri, George Koshy, Shondipon Laha, Steven Laird, Susan Larkin, Tamas Leiner, Patrick Lillie, James Limb, Vanessa Linnett, Jeff Little, Michael MacMahon, Emily MacNaughton, Ravish Mankregod, Huw Masson, Elijah Matovu, Katherine McCullough, Ruth McEwen, Manjula Meda, Gary Mills, Jane Minton, Mariyam Mirfenderesky, Kavya Mohandas, Quen Mok, James Moon, Elinoor Moore, Patrick Morgan, Craig Morris, Katherine Mortimore, Samuel Moses, Mbiye Mpenge, Rohinton Mulla, Michael Murphy, Megan Nagel, Thapas Nagarajan, Mark Nelson, Igor Otahal, Mark Pais, Selva Panchatsharam, Hassan Paraiso, Brij Patel, Natalie Pattison, Justin Pepperell, Mark Peters, Mandeep Phull, Stefania Pintus, Jagtur Singh Pooni, Frank Post, David Price, Rachel Prout, Nikolas Rae, Henrik Reschreiter, Tim Reynolds, Neil Richardson, Mark Roberts, Devender Roberts, Alistair Rose, Guy Rousseau, Brendan Ryan, Taranprit Saluja, Aarti Shah, Prad Shanmuga, Anil Sharma, Anna Shawcross, Jeremy Sizer, Manu Shankar-Hari, Richard Smith, Catherine Snelson, Nick Spittle, Nikki Staines, Tom Stambach, Richard Stewart, Pradeep Subudhi, Tamas Szakmany, Kate Tatham, Jo Thomas, Chris Thompson, Robert Thompson, Ascanio Tridente, Darell Tupper-Carey, Mary Twagira, Andrew Ustianowski, Nick Vallotton, Lisa Vincent-Smith, Shico Visuvanathan, Alan Vuylsteke, Sam Waddy, Rachel Wake, Andrew Walden, Ingeborg Welters, Tony Whitehouse, Paul Whittaker, Ashley Whittington, Meme Wijesinghe, Martin Williams, Lawrence Wilson, Sarah Wilson, Stephen Winchester, Martin Wiselka, Adam Wolverson, Daniel G. Wooton, Andrew Workman, Bryan Yates, and Peter Young.

